# Environmental Enrichment Reduces *adgrl3.1*-Related Anxiety and Attention Deficits but Not Impulsivity

**DOI:** 10.1101/2024.09.25.615074

**Authors:** Barbara D. Fontana, William H.J. Norton, Matthew O. Parker

## Abstract

Environmental factors play a role in the development and severity of neuropsychiatric disorders. Externalizing disorders (EDs) are linked to genetic and environmental factors and frequently co-occur with internalizing disorders (ID), such as anxiety. Individuals experiencing both ED/ID are often among the most likely to seek healthcare services, as this co-occurrence is associated with more severe symptomatology and greater functional impairment. Here, we investigate the role of environmental enrichment (EE) on a gene associated with impulsivity and attention deficits in zebrafish, *adgrl3.1*. Zebrafish were reared in either standard or enriched environments (from 15 days-post fertilization), and attention, impulsivity, and anxiety-related phenotypes were assessed at adult stages (4 months-post fertilization) using the open field test and 5-choice serial reaction time task. EE mitigated anxiety-related behaviors in *adgrl3.1* knockouts, normalizing locomotor patterns and decreasing thigmotaxis. Although attention deficits were reduced in *adgrl3.1*^*-/-*^ fish reared in EE, impulsive behaviors were not. Together, these findings suggest that while EE can alleviate some externalizing and internalizing phenotypes in zebrafish, impulsivity remains resistant to environmental modification. This study suggests that impulsivity is a stable trait in *adgrl3.1*^*-/-*^ fish, but highlights the importance of EE in mitigating some externalizing and internalizing symptoms.

**Summary statement:** Environmental enrichment reduces attention deficits and anxiety-like behavior, but not impulsivity in *adgr3.1* knockout zebrafish, highlighting the interaction between genetic and environmental factors in shaping externalizing and internalizing phenotypes.

## Introduction

Both ‘nature’ and ‘nurture’ play a role in the likelihood of developing neurodevelopmental and neuropsychiatric disorders. For example, genetic factors can explain a significant proportion of the variance in a wide range of psychiatric disorders (Wu et al., 2020), and aversive environments and/or chronic stress can increase both the relative odds of developing disorders (Schmitt et al., 2014) and symptom severity (Nugent et al., 2011). Supportive social environments and environmental enrichment (EE) have been shown to reduce stress and enhance cognitive function in both humans (Babin and Boles, 1996; Whitehouse et al., 2001) and animal models including rodents (Chauvet et al., 2009; Laviola et al., 2004) and zebrafish (Collymore et al., 2015; Marcon et al., 2018). Externalizing disorders (EDs) such as attention-deficit hyperactivity disorder (ADHD), as well as having a well-established genetic component (Barr and Dick, 2020), are significantly affected by the environment in which the child develops (Pressman et al., 2006; Schroeder and Kelley, 2009). In addition, a supportive and structured environment can foster greater social inclusion and improve learning in patients with psychiatric disorders (Halperin and Healey, 2011). Internalizing disorders (ID; *e.g*., anxiety and depression), which are commonly observed in patients with more extreme or treatment-resistant EDs (Knappe et al., 2022), can be partially alleviated when patients are part of an inclusive and supportive environment (Karawekpanyawong et al., 2021). Similar effects are observed in animal models exposed to EE, where it can enhance stress resilience and reduce internalizing symptoms like anxiety (Gatto et al., 2022; Lehmann and Herkenham, 2011; Vanisree and Thamizhoviya, 2021).

In animal models, the interactions between genotype and the environment can also modulate the severity of different psychiatric and neurological disorders (Emberti Gialloreti et al., 2019; Grizenko et al., 2008; Ronald et al., 2010). Genetic linkage studies have identified an ultra-conserved enhancer in *ADGRL3* that contributes to ADHD susceptibility by modulating gene expression, with noncoding variants affecting enhancer function and disrupting transcription factor binding (Arcos-Burgos et al., 2010; Bruxel et al., 2021; Martinez et al., 2016). *ADGRL3* variants are strongly associated with externalizing behaviors in animal models. For instance, our previous work on zebrafish harboring a knockout of *adgrl3.1* (the zebrafish homolog of *ADGRL3*) has revealed multiple changes to behaviors that can be used to model EDs, including hyperactivity, impulsivity, and attention deficits (Fontana et al., 2023; Lange et al., 2018; Lange et al., 2012; Sveinsdóttir et al., 2023). However, what is not presently understood is the role of the environment in the development of the behavioral phenotypes observed in this line.

Laboratory housing systems for zebrafish are typically designed based on economic and ergonomic aspects (Council, 2011), largely ignoring the animals’ natural habitat (*i.e*., well vegetated, moving water (Engeszer et al., 2007)). EE can mitigate the lack of concordance with the natural environment, and indeed, enrichment with plants and gravel substrate decreases stress-reactivity and anxiety in zebrafish (Marcon et al., 2018; Schroeder et al., 2014). EE has been studied in multiple species and, much like a supportive environment in humans (Farah et al., 2008), can alleviate symptoms of psychiatric disorders (DePasquale et al., 2016). Here, for the first time, we investigate the interactions between the *adgrl3.1* genotype and development in an enriched environment in zebrafish by adding artificial plants and gravel from developmental stages to adulthood on ED-related phenotypes, including hyperactivity, attention and impulsive behavior. We also studied the impact on anxiety-like behavior, an internalizing symptom often found increased on patients with externalizing disorders.

## Material and methods

### Animal husbandry, procedures and data reliability

Homozygous zebrafish (*adgrl3.1*^*-/-*^) generated by CRISPR-Cas9 as previously described (Sveinsdóttir et al., 2023) were bred in-house and kept in a re-circulating system on a 14/10-hour light/dark cycle (lights on at 9:00 a.m.), pH 8.4, at 28 °C (±1 °C). Zebrafish were fed with flake food (ZM-flake and ZM-300, ZM Ltd.) three times a day and adult brine shrimp during the mornings. To evaluate if EE affects attention deficits, impulsivity and anxiety-related patterns in *adgrl3.1*^*-/-*^, animals were split into groups according to two factors, 1) genotype (*adgrl3.1*^*-/-*^ *vs*. WT) and 2) environment (no-enrichment *vs*. EE). EE was added to the re-circulating system at 15 dpf. EE included gravel substrate and artificial leaves/grass as previously described (Lee et al., 2019). The non-enriched group were reared in the same re-circulating system, but in standard laboratory conditions which included an image of a gravel bed on the bottom of the tank but no substrate in the tank. Behavioral protocols included the 5-CSRTT, a continuous performance test, used to evaluate sustained attention and impulsivity (Everitt et al., 1983), and the open field tank, both of which are described in detail in the **Supplementary Materials**. At adult stages (4 months), zebrafish were tested in the open field task (*n =* 48) and then moved to a pair-housed system, with or without the addition of enrichment depending on the group, for the training of the 5-CSRTT (*n* = 48) for 6 weeks (**Fig. 1**). The number of animals per group were calculated a priori (d = 0.45, power = 0.65, alpha = 0.05) following extensive published work from our laboratory using zebrafish as a translational model (Alnassar et al., 2023; Fontana et al., 2020; Fontana et al., 2023) and considering our primary outcomes which resulted in a sample size of 48 animals. For the 5-CSRTT final sample size was smaller owing to the training demands of the test: 3 animals from each of the WT groups and 2 animals from *adgrl3.1*^*-/-*^ did not reach minimum criteria during training stages to proceed with testing analysis. All behavioral tests were performed from 10:00 to 15:00 and analyzed using the Zantiks AD system (Zantiks, Cambridge, UK). After 5-CSRTT, fish were euthanized using 2-phenoxyethanol from Aqua-Sed (Aqua-Sed™, Vetark, Winchester, UK).

**Fig. 1.**
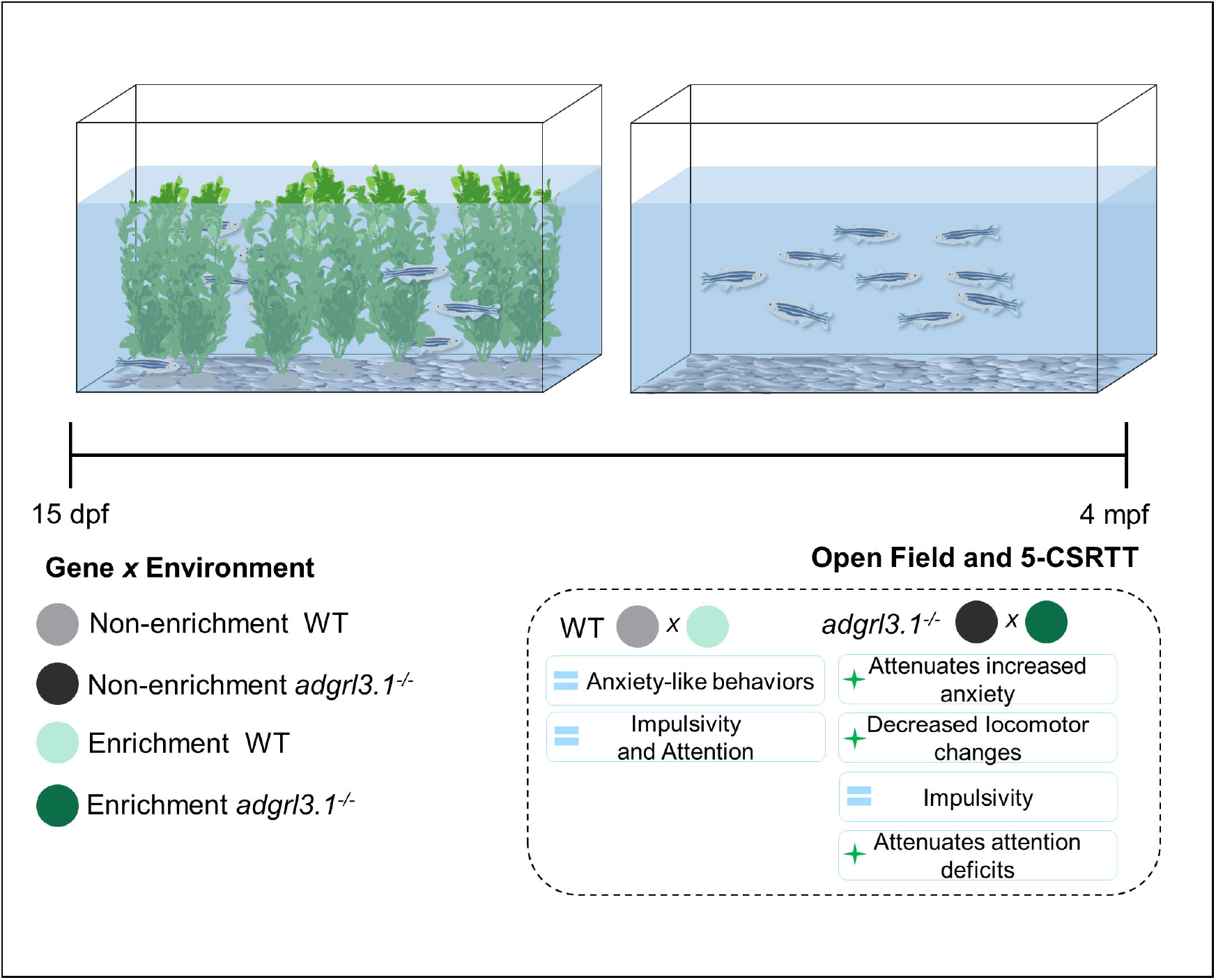
Illustration of the experimental design showing the time in which animals are transferred to environmental enrichment (15 days-post fertilization – dpf) and when zebrafish are tested for behavior (4 months-post fertilization – mpf). Main findings are represented in the dotted box, showing that environmental enrichment did not affect WT animals, but influenced *adgrl3.1* anxiety-like behaviors and attention deficits.

Data was collected from two independent experimental batches to ensure data reliability. Animal Welfare and Ethical Review Board, and under license from the UK Home Office (Animals (Scientific Procedures) Act, 1986) [PPL: P9D87106F].

### Statistics

Data were analyzed using GraphPad Prism and the results were expressed as mean ± standard error of the mean (S.E.M). Normality was assessed prior to testing. Two-way ANOVA was used to evaluate the effects of genotype (two levels – WT *vs. adgrl3.1*) *vs*. environment (two levels – non-enriched *vs*. EE) in the open field and in the 5-CSRTT. Tukey’s test was used as a post-hoc analysis. Results were considered significant when p ≤ 0.05.

## Results

### Environmental enrichment decreases alterations in locomotion and thigmotaxis of adgrl3.1 knockouts

**Fig. 2** depicts the interactions between genotype (*adgrl3.1*^*-/-*^ vs WT) and environment (enriched *vs*. non-enriched). For locomotion, there was a significant main effect of genotype, with *adgrl3.1*^*-/-*^ more hyperactive (F _(1, 44)_ = 4.393; *p*=* 0.0419) and a significant genotype*environment interaction (F _(1, 44)_ = 5.392; *p*=* 0.0249). No main effect of environment was observed (F _(1, 44)_ = 1.1441; *p =* 0.2364). The decrease in locomotion in the *adgrl3.1*^*-/-*^ was reversed in the enrichment condition compared to the non-enriched. When looking at thigmotaxis (time spent close to the walls), while no main effects for EE (F _(1, 44)_ = 3.835; *p =* 0.0566) or genotype (F _(1, 44)_ = 0.78998; *p =* 0.3790) was observed, a significant effect for the interaction between environment and *adgrl3.1* genotype (F _(1, 44)_ = 6.180; *p* =* 0.0168) was found. This was characterized as a significant increase in thigmotaxis for *adgrl3.1*^*-/-*^ compared to WT controls (non-enriched) (*p\** = 0.0153). When grown in EE, *adgrl3.1*^-/-^ did not show differences in thigmotaxis behavior compared to controls (*p* = 0.8735).

**Fig. 2.**
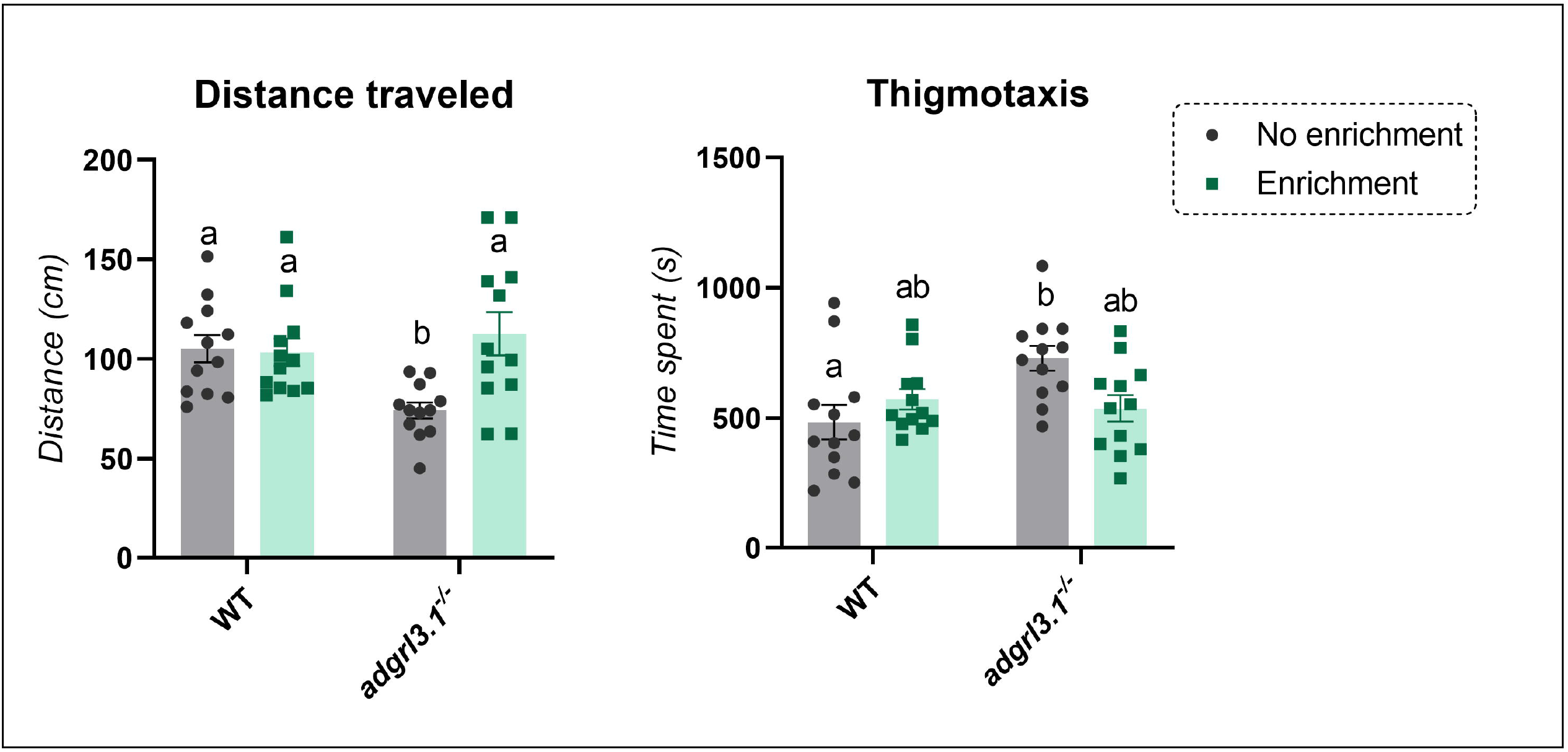
Locomotion and anxiety-like behavior of WT vs. *adgrl3.1*^*-/-*^ grown in different environments in the open field tank. Tukey’s test was used as post-hoc analysis and different letters indicate significant statistical difference (*p* < 0.05; *n* = 12). The data is represented as mean ± S.E.M.

### Attention deficits are reduced in adgrl3.1 knockout fish that grew in an enrichment environment

Regarding attention and impulsive behavior, a significant effect of genotype effect was observed for accuracy (F _(1, 33)_ = 8.883; *p** =* 0.0054) with no interaction (environment *vs. adgrl3.1* genotype; F _(1, 33)_ = 0.1694; *p =* 0.6833) or environment effect (F _(1, 33)_ = 1.933; *p =* 0.1737). Tukey’s post-hoc analysis showed a significant effect for WT non-enriched and enriched *vs. adgrl3.1* non-enriched (*p* < 0.05). However, with respect to impulsive behavior, there was a significant effect of genotype (F _(1, 33)_ = 39.58; *p**** <* 0.0001) with an increase of responses for both *adgrl3.1* groups compared to WT animals, and this was not affected by environment (*p* < 0.05). No significance was detected for interaction (environment *vs. adgrl3.1* genotype; F _(1, 33)_ = 1.442; *p =* 0.2382) or environment (F _(1, 33)_ = 0.04303; *p =* 0.8369) in the anticipatory responses. No significant interaction (environment *vs. adgrl3.1* genotype; F _(1, 33)_ = 0.09278; *p =* 0.7626), environment (F _(1, 33)_ = 0.08638; *p =* 0.9265) or *adgrl3.1* genotype (F _(1, 33)_ = 0.5973; *p =* 0.4451) effect was observed for the omissions (**Fig. 3**).

**Fig. 3.**
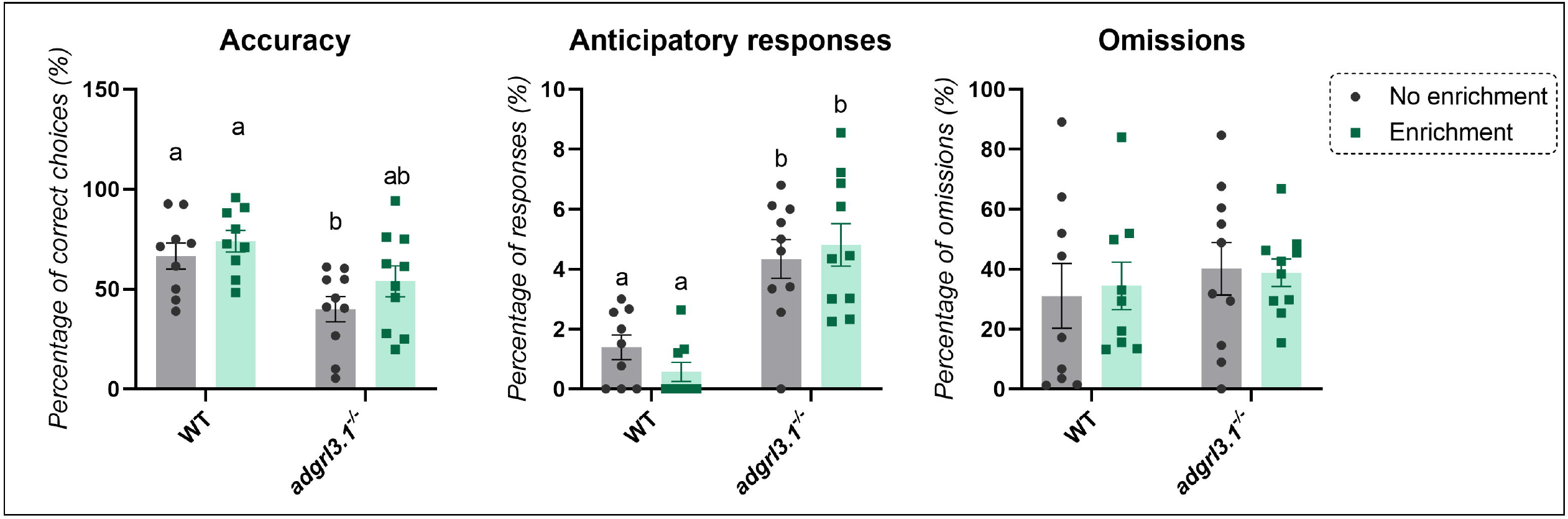
Behavioral response of WT vs. *adgrl3.1*^*-/-*^ grown in different environments in the 5-CSRTT. Tukey’s test was used as post-hoc analysis and different letters indicate significant statistical difference (*p* < 0.05; *n* = 9 – 10). The data is represented as mean ± S.E.M.

## Discussion

We have shown, for the first time, that EE can protect against some internalizing and externalizing phenotypes observed in *adgrl3/1*^*-/-*^ fish but, critically, not all. We found that deficits in attention and anxiety were attenuated by EE; however, impulsivity was not affected, with the impulsive behavioral phenotype the same across different contexts. These findings suggest that EE can have a positive role on decreasing some externalizing and internalizing behaviors but are limited by behavioral domain. Impulsivity is a stable behavioral trait in *adgrl3.1*^*-/-*^ fish and appears to be impervious to the beneficial effects of EE.

Environmental factors in humans, such as behavioral intervention (cognitive stimulation and physical activity) and lifestyle (frequency of cognitive and social activities) have shown important benefits on patients with dementia and brain injuries (Sampedro-Piquero and Begega, 2017). In zebrafish, EE has positive effects on anxiety and cognition, being able to decrease anxiety-like behavior similarly to anxiolytic drugs (Giacomini et al., 2016b). For example, EE (DePasquale et al., 2016) and anxiolytic drugs, such as diazepam and fluoxetine (Gebauer et al., 2011; Giacomini et al., 2016a), reduce animals time spent in the bottom in the novel tank diving task, suggesting that EE can be used as an approach to reduce stress in those fish.

Here, we used EE to evaluate the role of a positive environment for *adgrl3.1* knockout. Although the core symptoms of externalizing disorders such as ADHD involve attention deficits, impulsivity and hyperactivity, these disorders are often comorbid with anxiety disorders (Blazer, 1982; Fayyad et al., 2017; Heim et al., 2008; Levy et al., 2020). We have previously shown that *adgrl3.1* knockout zebrafish show increased anxiety-like behavior and that abnormal behaviors can be worsened by external factors such as social isolation (Fontana et al., 2024). Here, we used an additional test to confirm that *adgrl3.1* knockouts display increased anxiety-like behavior by increasing thigmotaxis (time spent close to the wall), and EE was able to attenuate the role of this gene on this behavior. Importantly, locomotion was decreased in the knockout fish, which could be caused by either decreased exploration of the tank or increased immobility. Although these parameters have not been assessed before, similar findings were previously observed on *adgrl3.1* knockouts, where hyperactive patterns are only observed after adaptation to a new environment and not when first introduced (Fontana et al., 2023). Interestingly, not only thigmotaxis but also distance traveled alterations were improved in the presence of EE. These data suggests that *adgrl3.1* knockouts initially show high anxiety in a new environment, and only after habituation do animals start to display a hyperactivity movement pattern. However, when these animals grow in an EE, the anxious response behaviors to a new environment are decreased, highlighting the importance of EE on internalizing symptoms.

The mechanisms in which EE may prevent the negative impact of disease in zebrafish are unknown. In rats, for example, EE is shown to significantly reduce anxiety in the open field test leading to decreased depressive-like behavior and increased serotonin concentrations in the prefrontal cortex (Brenes et al., 2008). Similarly, a study of gilthead seabream (*Sparus aurata*) showed that individuals reared in EE show reduced 5-HT and DA-related activity under higher social stress conditions (Batzina et al., 2014). Thus, decreased 5-HT and DA activity after acute stressor could be a possible mechanism underlying EE protector effects in a zebrafish model of externalizing disorders and this should be further explored.

EE attenuated the deficits in attention observed in *adgrl3.1* knockout animals. These findings support previous work using the Spontaneously Hyperactive Rat (SHR) model for ADHD, where EE improved the animals’ cognitive performance across a range of tasks, including open field habituation, water maze spatial navigation and a social and object recognition task (Pamplona et al., 2009). There is a clear literature on ADHD and EE highlighting the importance of the environment in clinical cases, where there are long-lasting positive effects on cognition and improvements in many aspects of ADHD, such as better insertion in social contexts and learning improvements (Halperin and Healey, 2011). Collectively, this supports the relevance of the environment interaction with ADHD-symptoms caused by alterations in the *ADGRL3* gene, specifically when looking at attention and anxiety-like behavior. Our observation that EE improves attention and reduces anxiety in *adgrl3.1*^*-/-*^ fish, but has no effect on impulsivity, may shed important light on the mechanisms of the genetic contribution to ADHD and other externalizing disorders. For example, it suggests that while certain behavioral aspects, such as attention and anxiety, can be more influenced by environmental factors, impulsivity may be more deeply rooted in genetic predispositions linked to the *adgrl3.1* gene.

In the context of ADHD, impulsivity is a core symptom that is often resistant to environmental interventions, as observed in our previous work where following social isolation *adgrl3.1* knockouts did not show changes in impulsivity in a novel seeking task (Fontana et al., 2024). The persistence of impulsivity despite EE implies that impulsivity in ADHD might be predominantly genetically determined and less susceptible to modification through lifestyle or environmental changes. This aligns with clinical observations where the combination of therapy and pharmacotherapy do not have a greater impact on impulsivity comparing to pharmacotherapy alone in individuals with ADHD (Corbisiero et al., 2018). Meanwhile, anxiety for example, is often well-managed with behavioral therapies (Cuijpers et al., 2013; Otto et al., 2006). Thus, while EE can improve some symptoms, the imperviousness of impulsivity in response to environmental change highlights the need for targeted therapeutic strategies, particularly those addressing the genetic pathways involved, to effectively manage these complex debilitating disorders.

Overall, we saw that environment can significantly influence internalizing behaviors such as anxiety-related phenotypes as well as altering the locomotion of *adgrl3.1* knockout, an animal model for externalizing disorders. Although impulsivity was still higher in *adgrl3.1* knockouts that grow in a enriched environment, the attention of *adgrl3.1* knockouts compared to controls was improved by enrichment. These data suggest that an enriched environment can positively modulate some of the behavioral alterations displayed by *adgrl3.1* knockouts. Altogether, these data will be useful in better understanding how environmental factors influence the severity of externalizing and internalizing disorders, and how positive environments may protect against behavioral alterations associated with genetic changes, such as those associated with the *adgrl3.1* gene.

## Author contributions

Conceptualization: B.D.F., M.O.P.; Methodology: B.D.F.; Formal analysis: B.D.F.; Investigation: B.D.F.; Resources: W.H.J.N., M.O.P; Data curation: B.D.F.; Writing - original draft: B.D.F.; Writing - review & editing: W.H.J.N., M.O.P; Visualization: B.D.F.; Supervision: W.H.J.N., M.O.P; Funding acquisition: W.H.J.N., M.O.P.

## Data and resource availability

All data is fully available on a GitHub repository (github.com/BarbaraDFontana/Enrichment_adgrl3.1_zebrafish).

