## Supplementary Material for "Environmental Enrichment Reduces *adgrl3.1*-Related Anxiety and Attention Deficits but Not Impulsivity"

**Supplementary Methods**

*Open Field Test*

The open field test is a behavioural test commonly used for measuring locomotion, exploratory activity and anxiety-like behaviour in adult zebrafish (Liu et al., 2018; Stewart et al., 2012). Animals were placed individually in the open field (20 cm length x 15 cm width, 10 cm water column depth) and behaviour was recorded for 20 minutes. All behaviours were recorded and analysed using Zantiks AD system (Zantiks Ltd., Cambridge, UK). The tank was separated in two virtual areas (central and peripheral area, 2 cm close to the wall) to provide a detailed evaluation of exploratory activity. Lighting was controlled for each tank size and range from 290 to 343 LUX, and the water was changed between each individual. The following endpoints were measured: distance travelled (cm) and time (s) spent in the peripheral zone (thigmotaxis).

*5-CSRTT training phases and testing*

The 5-CSRTT is a continuous performance test often used to evaluate sustained attention and impulsivity (Everitt et al., 1983). For this task, zebrafish are trained to respond to a light stimulus in one of five spatially distinct locations at the back of the test apparatus. Impulsivity is ascertained by examining the animal’s ability to withhold its response to the light stimuli during a pre-stimulus interval. This behavioral test have been previously validated pharmacologically for zebrafish aiming to assess impulsivity when the animal responds by approaching the stimulus before the onset of a light (Parker et al., 2014; Parker et al., 2013a; Parker et al., 2012). The test is performed in five steps as described below.

*1. Light stimulus training:* Fish were trained for five days to trigger a response from the initiator and the five stimulus lights, allowing them to habituate to the light stimuli (one session of 30 trials/day). All lights were illuminated simultaneously including the initiator area and the five openings at the back of tank. The fish was required to enter in any of light areas to receive food as a reward stimulus. If animals do not respond in 60 seconds (limited hold), no food reward is delivered, and a new test begins after an interval of 20 seconds.

*2. Initiator Light Training:* During this second stage, fish learned to activate the initiator light (illuminated for 60 seconds). During this phase, when animals swim to the initiator light, it illuminates the feed hopper light and a food reward is delivered. In this phase, animals were trained for 5 days (one session of 30-trials/day) until animals reached an average of initiator light activation of 80% from 30 trials.

*3. Stimulus Light Reward:* In this phase, fish learned to approach any of the five illuminated stimulus lights to receive a food reward after triggering the initiator light (beginning test). Animals underwent training for 5 days at this stage, achieving an average of 60% correct responses before the next step was initiated. At this stage, three animals from each WT group were removed and two animals from each *adgrl3.1* knockout groups were removed due to their poor performance on the test.

*4. Stimulus Light Discrimination:* Here, animals learned to discriminate between the individual stimulus lights. Once the fish activates the initiator light, only one of the stimulus lights was illuminated (5-seconds delay). In subsequent trials, individual stimulus lights were illuminated in a random order. The fish were required to approach the illuminated stimulus during a 50-sec illumination, and correct responses were rewarded with food delivery in the feed area. Animals were trained until they achieved a correct response rate of 60%, based on the animals’ total number of choices (5 days training).

*5. 5-CSRTT:* The final stage is similar to the light discrimination phase where, following triggering the initiator light, one of five stimulus lights were briefly illuminated (50-sec). Then, the fish were required to approach the correct light within the illumination time in order to receive a food reward. However, in this stage a 10-seconds delay between activation of initiator light and the five-stimulus light is added to assess animals’ anticipatory responses. The parameters evaluated in 5-CSRTT were: number of correct responses, number of incorrect responses, omissions (start the session without swimming to any of the 5-choice chambers), and premature responses (approach to one of the 5-choice chambers before the light stimulus). This test was performed during three consecutive days and the average response was used to ensure consistent performance on the task. These parameters enabled the analysis of impulsiveness (premature responses) and attention (correct responses) in adult zebrafish. The 5-CSRTT data was collected using the Zantiks AD system (Zantiks Ltd., Cambridge, UK).
